# Cardiac myocytes respond differentially and synergistically to matrix stiffness and topography

**DOI:** 10.1101/682930

**Authors:** William Wan, Kristen K. Bjorkman, Esther S. Choi, Amanda L. Panepento, Kristi S. Anseth, Leslie A. Leinwand

## Abstract

During cardiac disease progression, myocytes undergo molecular, functional and structural changes, including increases in cell size and shape, decreased myocyte alignment and contractility. The heart often increases extracellular matrix production and stiffness, which affect myocytes. The order and hierarchy of these events remain unclear as available *in vitro* cell culture systems do not adequately model both physiologic and pathologic environments. Traditional cell culture substrates are 5-6 orders of magnitude stiffer than even diseased native cardiac tissue. Studies that do account for substrate stiffness often do not consider intercellular alignment and *vice versa*. We developed a cardiac myocyte culture platform that better recapitulates native tissue stiffness while simultaneously introducing topographical cues that promote cellular alignment. We show that stiffness and topography impact myocyte molecular and functional properties. We used a spatiotemporally-tunable, photolabile hydrogel platform to generate a range of stiffness and micron-scale topographical patterns to guide neonatal rat ventricular myocyte morphology. Importantly, these substrate patterns were of subcellular dimensions to test whether cells would spontaneously respond to topographical cues rather than an imposed geometry. Cellular contractility was highest and the gene expression profile was most physiologic on gels with healthy cardiac tissue stiffness. Surprisingly, while elongated patterns in stiff gels yielded the greatest cellular alignment, the cells actually had more pathologic functional and molecular profiles. These results highlight that morphological measurements alone are not a surrogate for overall cellular health as many studies assume. In general, substrate stiffness and micropatterning synergistically affect cardiac myocyte phenotype to recreate physiologic and pathologic microenvironments.

**Significance Statement:** Heart disease is accompanied by organ- and cellular-level remodeling, and deconvoluting their interplay is complex. Cellular-level change is best studied *in vitro* due to greater control and uniformity of cell types compared to animals. One common metric is degree of cellular alignment as misalignment of myocytes is a hallmark of disease. However, most studies utilize featureless culture surfaces that are orders of magnitude stiffer than, and do not mimic the scaffolding of, the heart. We developed a hydrogel platform with tunable stiffness and patterns providing topographical alignment cues. We cultured heart cells on and characterized multifactorial responses to these dynamic surfaces. Interestingly, conditions that yielded greatest alignment did not yield the healthiest functional and molecular state. Thus, morphology alone is not an indicator of overall cellular health.

## Introduction

Cardiac myocytes undergo many molecular, biophysical, and biochemical changes during the progression of heart disease. These changes include myofibrillar and sarcomeric disarray, cell spreading and aspect ratio changes, interactions with extracellular matrix (ECM) proteins, expression of cell-cell interaction proteins, and alterations in gene expression (1–5). Myocytes interact with their extracellular microenvironment in a feedback loop that ultimately determines whether they undergo physiological or pathological remodeling that may culminate in changes in cell contractility. However, studying these changes *in vitro* is challenging, as traditional cell culture studies are conducted on smooth, isotropic surfaces made of tissue culture polystyrene (TCPS) or glass, whose stiffnesses are 5-6 orders of magnitude greater than that of native tissue. Moreover, in these systems, there is minimal control of the density, composition, and mechanics of the extracellular environment. As a result, researchers delving into the cell biology of heart disease have been unable to reproduce many of the complexities of the *in situ* environment of native hearts.

Motivated by this gap, advances in synthetic biomaterial substrates have allowed the creation of synthetic ECM mimics that enable precise control over the presentation of substrate stiffness (6, 7), topographical cues (8), presentation of ECM proteins, and the tethering and removal of growth factors (6). These biomatrices share many properties with tissues and allow one to perform genetic and biochemical manipulations of the cellular environment, while simultaneously controlling cell-matrix interactions. Poly(ethylene glycol) (PEG) is one commonly used biomaterial substrate, and PEG is often selected because of its biocompatibility, hydrophilicity, resistance to nonspecific protein adsorption, and ability to tune its mechanical properties to match various biological tissues. Furthermore, adhesive ligands and/or cytokines can be conjugated or released at user-defined points in time and space (6). These collective properties have rendered PEG hydrogels useful for conducting experiments that are largely intractable on static plastic or glass surfaces. Examples of such studies have been reviewed elsewhere (7–9). Here, an advanced PEG hydrogel system with light-tunable properties was used to investigate how neonatal rat ventricular myocytes (NRVMs) respond to a simultaneous presentation of substrate stiffness cues and patterns.

The stiffness (as assessed by Young’s modulus) of healthy neonatal rat myocardium is ∼4-11kPa, and the stiffness of healthy adult rat myocardium is ∼11-46kPa. The stiffness of infarcted areas in adult rats can reach up to 56kPa (10), while the stiffness of hearts with other fibrotic diseases ranges from ∼35-50kPa (10–13). In the native myocardium, cells are arranged in a parallel, “brick-wall” pattern, while in fibrotic disease, there are numerous changes in the alignment of cardiac myocytes and in the composition and orientation of ECM proteins (14). Other groups culturing cardiac myocytes on hydrogels and flexible surfaces have demonstrated that cardiac myocytes exhibit greatest striation and maximum work and contractile force on substrates with stiffnesses between 10-17kPa, which closely matches the stiffness measured in neonatal and embryonic hearts (15, 16). Culturing cardiac myocytes on soft hydrogel substrates also attenuates the expression of pathological (fetal) genes such as *Nkx2.5* and *Anf* (*Nppa*) (17, 18).

Beyond control of the matrix mechanical properties, techniques such as molding and microcontact printing have been used to recreate the patterned arrangement and aspect ratio of cells seen in the healthy myocardium (19–25). As expected, cell aspect ratio and alignment are both greater when NRVMs are cultured on patterned substrates than on smooth surfaces (20, 26). Specifically, culturing NRVMs on substrates micropatterned with topographical features of parallel 20 μm-wide fibronectin lines (27), 20 μm-wide eroded channels (22), or printed with high aspect ratio rectangular adhesion 2000-2500 μm^2^ islands (28, 29) leads to higher levels of sarcomeric alignment than on isotropic surfaces. From a functional perspective, NRVMs cultured on patterned substrates generate greater peak systolic stress than cells cultured on isotropic substrates (20, 27). In a system where NRVMs were cultured on thin, flexible 80×12 μm polydimethylsiloxane (PDMS) strips arrayed in a brick wall pattern, McCain *et al.* found increased α myosin heavy chain (αMHC; *Myh6*)-to-βMHC (βMHC; *Myh7*) ratios and on patterned surfaces vs. isotropic surfaces (20). Because lower αMHC-to-βMHC ratios are correlated with cardiac disease, the results of this study suggest that patterning matrix cues may promote a healthier NRVM phenotype. However, aspect ratio *alone* cannot always predict whether a cardiomyocyte is in a physiologic or pathologic state (reviewed in (1, 30). For example, the aspect ratio of a typical adult cardiomyocyte is approximately 7:1. Pressure overload due to pathologic stimuli such as high blood pressure or aortic valve stenosis, as well as physiologic stimuli such as resistance weight training can both result in aspect ratios less than 7. By contrast, volume overload due to pathologic stimuli such as valve regurgitation, ventricular septal defects, or myocardial infarctions as well as physiologic stimuli such as running or swimming can both result in aspect ratios greater than 7. Despite this wealth of experimental observations, there remains a paucity of information as to how extracellular signals influence intracellular signaling in NRVMs, and very few literature reports integrate an in-depth biomolecular characterization of NRVMs while simultaneously controlling and presenting multiple substrate microenvironmental cues.

In recent years, a number of studies have revealed that constraining cardiomyocytes to different dimensions can result in changes in morphological and functional properties. The current study described here builds from this foundational work by examining whether micropatterned hydrogel substrates with feature dimensions smaller than an individual cell (ensuring cells remain on top of the features) are sufficient to serve as simple topographical cues to modulate molecular, structural, and functional cell-autonomous and intercellular properties. Such changes can be interpreted as integration and interpretation of environmental mechanical signals rather than a response to a forced morphological constraint. We further examine the intersection of substrate topography and stiffness to evaluate the possibility of modeling physiologic and pathologic conditions with one tunable culture platform.

## Results

### NRVMs form gap junctions and contract on micropatterned photoresponsive hydrogels

Patterning was performed on photoresponsive PEG hydrogels for culturing NRVMs and the process is schematically shown in Figure 1 and previously described in (31). Substrates were formulated to have a stiffness of that of either a healthy neonatal heart (10kPa) or a diseased heart (35kPa). Regular, rectangular micropatterns of dimensions 40×5µm, 20×5µm, 10×5µm, and 5×5µm were formed on the surface via photodegradation. The rectangular features were spaced 5µm apart, and typically, the NRVMs spread across many of the topographical features. The feature sizes resulted in patterns with aspect ratios ranging from 1:1 to ∞:1 (Figure 1). NRVMs in all culture conditions, including TCPS, spontaneously contracted and formed mature sarcomeres (Figure 2: representative images of TCPS and 10kPa ∞:1), but sarcomeres in cells on hydrogels were better organized as revealed by F-actin staining. Collectively, these results suggest that, similar to culturing on TCPS, NRVMs cultured on both smooth and patterned PEG substrates maintained a cardiac phenotype measurable at the protein and functional levels with respect to sarcomere structure and spontaneous contractions.

**Figure 1.**
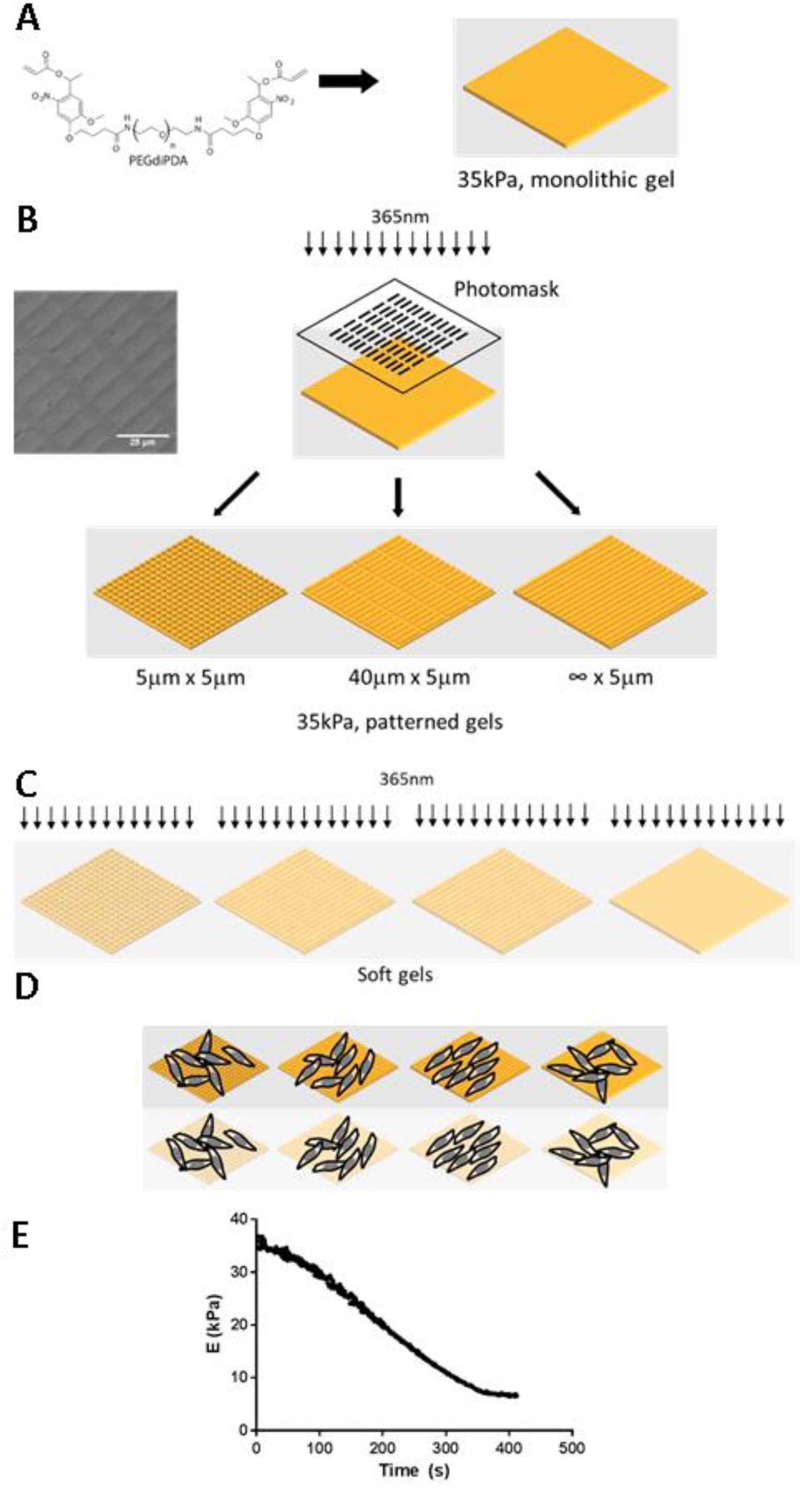
Fabrication and characterization of smooth or stiff photopatterned PEG hydrogel cell culture substrates. **A.** A monolithic gel with Young’s stiffness of 35kPa was fabricated with entrapped gelatin as an adhesive ECM ligand. Microtopographies were generated by directing collimated 365nm light (15mW/cm^2^) through a photomask. Feature dimensions were 5µm × 5µm, 10µm × 5µm, 20µm × 5µm, and 40µm × 5µm. Channels were 5µm wide. All features were spaced 5µm apart. **B.** Gels were then softened to 10kPa by irradiating with 365nm (10mW/cm^2^) light to decrease crosslinking density at the gel surface. **C.** To generate soft substrates, gels with an initial stiffness of 35kPa were irradiated for 5 minutes to reach a final soft stiffness of 10kPa. **D.** After swelling in PBS for 2-3 days, NRVMs were seeded on the patterned gels at a density of 50,000 cells/cm^2^. **E.** Time-sweep of the Young’s modulus when the hydrogel was exposed to 365nm (10mW/cm^2^) light.

**Figure 2.**
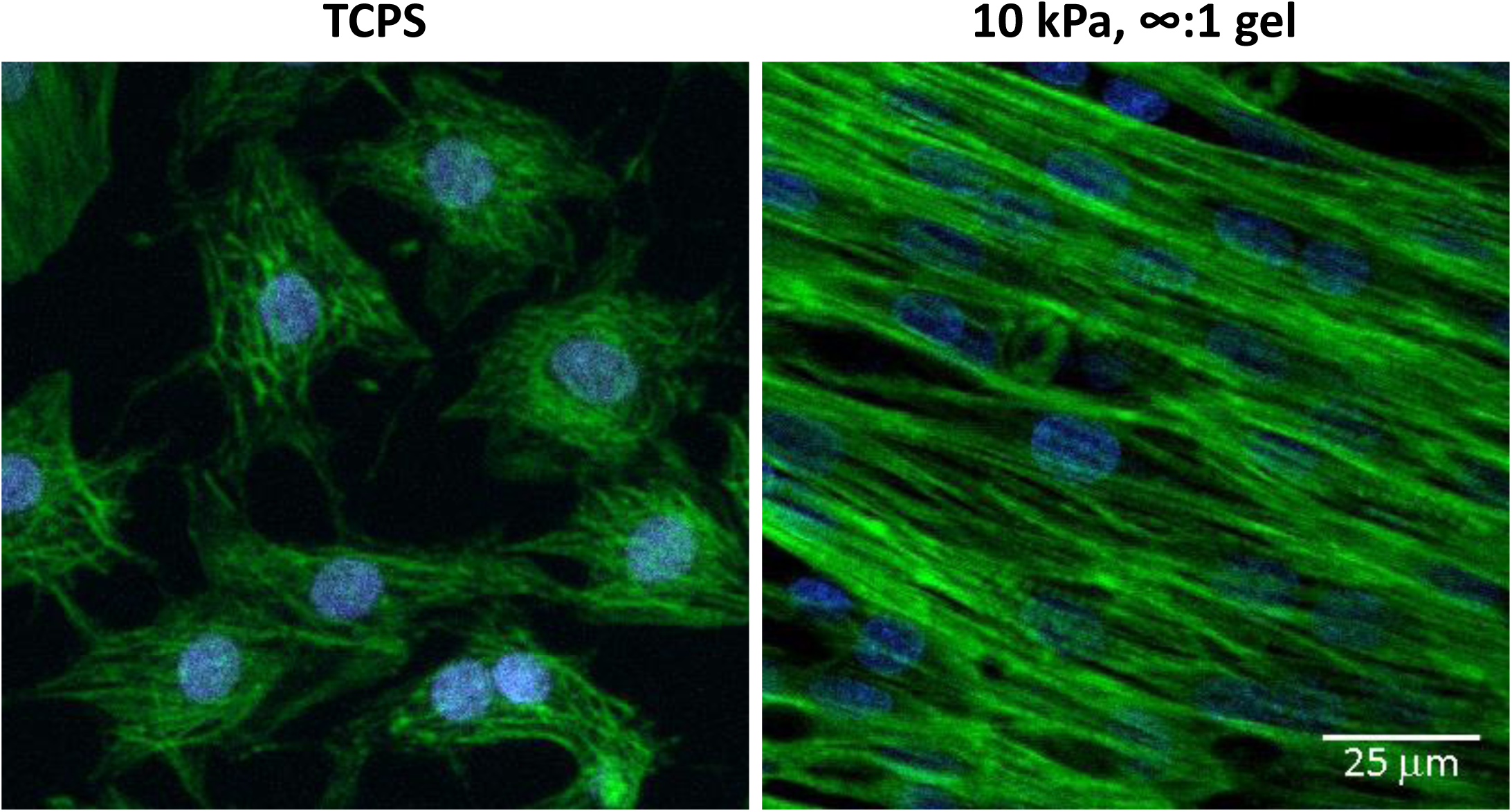
Sarcomeres in cells cultured on hydrogels are more organized than cells cultured on TCPS. Fluorescent staining of F-actin (green) in NRVMs cultured on TCPS or 10kPa gels patterned with 5 µm channels (∞:1 aspect ratio). Nuclei were stained with DAPI (blue). NRVMs were also spontaneously contracting by day 4 of culture.

### NRVM and F-actin alignment is proportional to the pattern aspect ratio

Higher aspect ratio patterns (i.e., from squares to channels) resulted in higher levels of cell alignment in the direction of the pattern and lower variation in the cell orientation. The major axis of the cell nuclei relative to the patterns was used to calculate the distribution of orientational angles (Figure 3A,B). The circular standard deviation of the angular differences measures the degree of cell alignment, where low angular standard deviations represent a high degree of cell alignment and high angular standard deviations represent more random alignment of cells (Figure 3C). Infinitely long channels produced the largest number of NRVMs that were perfectly aligned with the pattern, while smooth gels and 1:1 patterns resulted in cells that were more randomly oriented (Figure 3B). Of further note, NRVMs cultured on stiffer gels were more closely aligned to the pattern compared to those on softer matrices with the same pattern. Fold changes in percent alignment observed here were consistent with changes in myocyte alignment in studies of wildtype mice (∼46% aligned) and mice with hypertrophic cardiomyopathy (∼25% aligned) (32).

**Figure 3.**
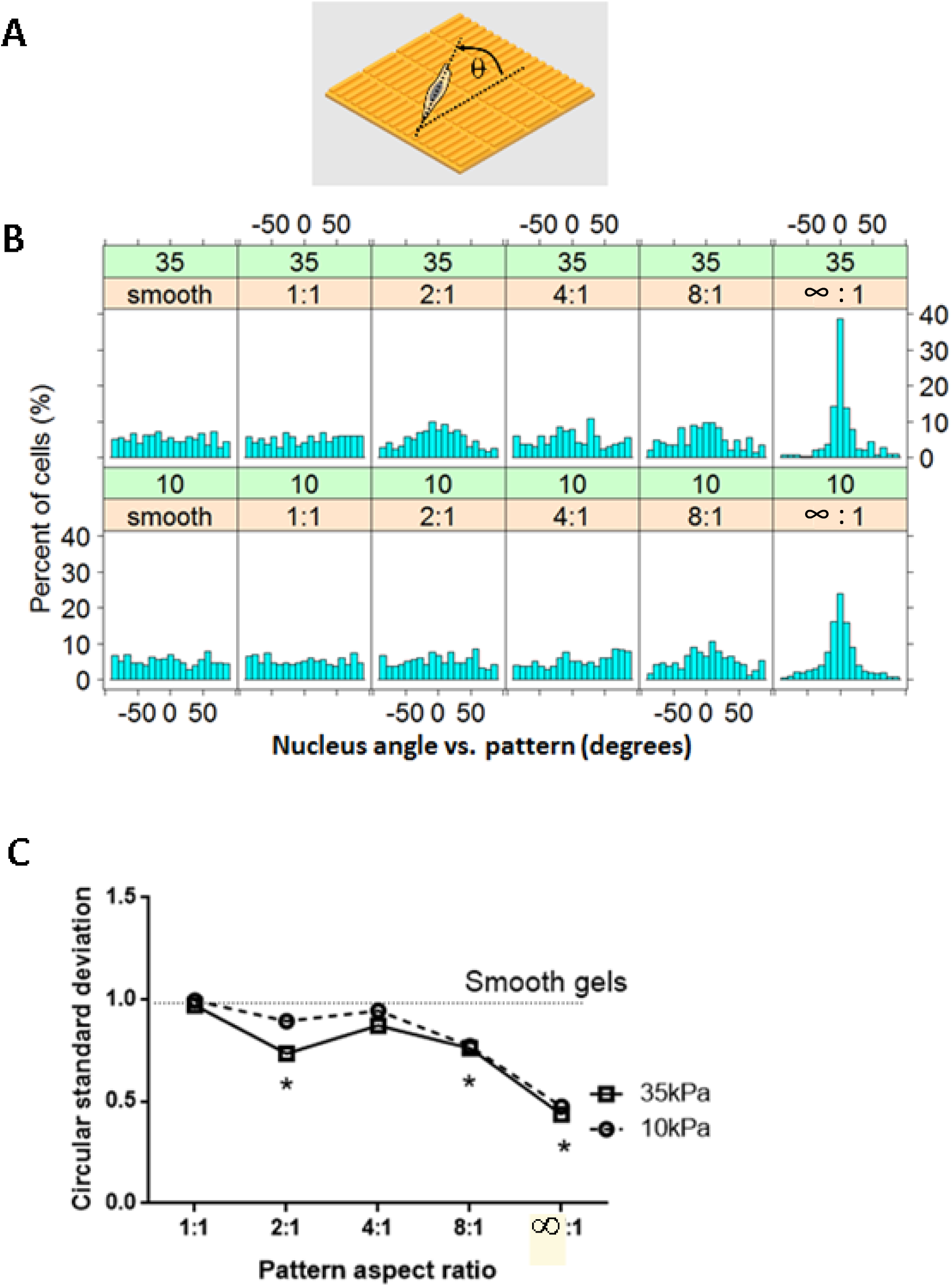
Cells exhibit greater alignment to the substrate pattern as the aspect ratio of the pattern increases from 1:1 to ∞:1. **A.** The angular difference between the pattern and the major axis of each cell nucleus was calculated. (not to scale) **B.** The horizontal axis represents the angular difference between the substrate major axis and the major axis of cell nuclei. **C.** The increase in nuclear alignment with the pattern can be visualized by plotting the variance of the histograms from the above panes. On 8:1 and ∞:1 patterns, the circular standard deviation was significantly lower than smooth gel and TCPS conditions for both 10kPa and 35kPa gels. n = 200-400 cells per group. * indicate p < 0.05 vs. smooth group based on the equal kappa test.

Cell aspect ratio and F-actin alignment also increased in a manner that correlated with the pattern aspect ratio (Figure 4A). Cell elongation, as measured by the aspect ratio, increased as the pattern aspect ratio went from 1:1 to ∞:1 on both 10kPa and 35kPa gels. Regression slopes, representing the relationship between pattern and cell aspect ratios, were statistically significant for both soft and stiff gels (Figure 4B, Table 1). NRVMs cultured on smooth substrates adopted a slightly lower aspect ratio, as did NRVMs cultured on 1:1 soft gels (Figure 4B). On rectangular patterns with aspect ratios of 2:1 or greater, the aspect ratio for cells on gels was significantly greater than on TCPS; however, there were no significant differences in cell aspect ratio between soft and stiff gels. These results suggest that, under the experimental conditions tested, the aspect ratio of the underlying pattern had a greater impact on cell aspect ratio than substrate elasticity.

**Table 1:**
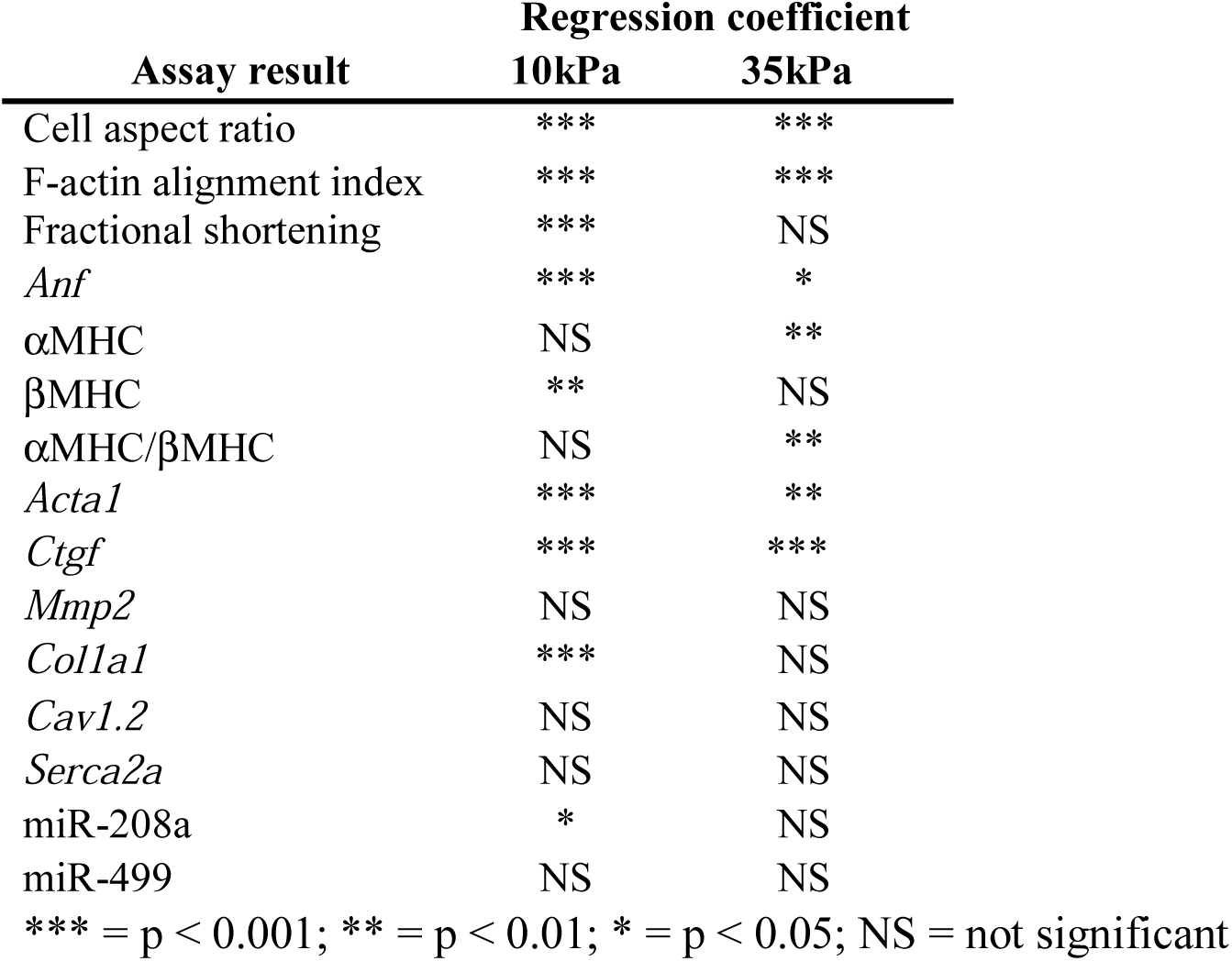
Significance levels (p-values) of regression slopes.

**Figure 4.**
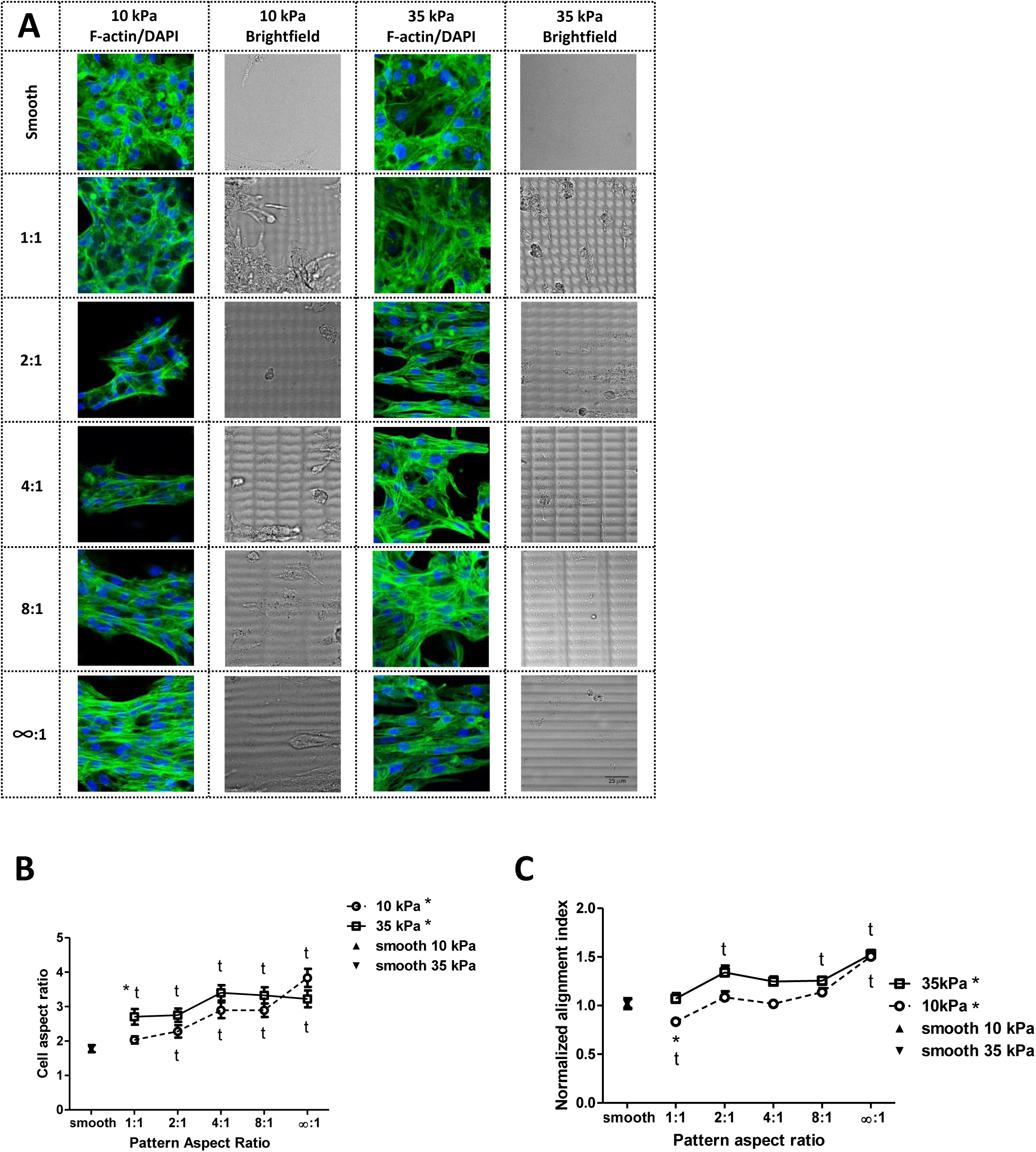
Topographical cues direct NRVM morphology and F-actin alignment. **A.**Fluorescent and brightfield images show greater F-actin alignment within cells for NRVMs cultured on substrates with increasing pattern aspect ratio. A relatively cell-free area neighboring the region chosen for the F-actin/DAPI panel in the same field of view was selected to reveal the gel pattern. **B.** Cell aspect ratios increase with increasing pattern aspect ratio. **C.** F-actin alignment increases with increasing pattern aspect ratio and is greater in 35kPa gels than in 10kPa gels for 1:1 aspect ratio gels. * next to legend indicates p < 0.05 for regression slope * = p < 0.05 for 10kPa vs. 35kPa, t = p < 0.05 vs. TCPS

F-actin alignment, a measure of internal sarcomeric organization, also increased with increasing pattern aspect ratio. A fast Fourier transform (FFT) algorithm was used to quantify F-actin alignment in NRVMs cultured on both soft and stiff patterned substrates. F-actin alignment on hydrogels was normalized to the F-actin alignment of NRVMs on TCPS. Regression slopes were significant for both 10kPa and 35kPa gels, indicating a significant correlation between the underlying pattern aspect ratio and F-actin alignment (Table 1). F-actin alignment on soft gels was not significantly different than on TCPS for intermediate pattern aspect ratios of 2:1 through 8:1 (Figure 4C). On channeled substrates (∞:1), the percent F-actin alignment was higher than on TCPS and was nearly identical on soft and stiff substrates. On 1:1 aspect ratio gels, F-actin alignment was significantly higher on stiff gels.

### Fractional shortening increases with pattern aspect ratio on soft, but not stiff, gels

NRVM function was assessed by measuring contractility through fractional shortening. On soft gels, there was a significant, positive relationship between pattern aspect ratio and fractional shortening (Table 1). For aspect ratios greater than 2:1, fractional shortening was also significantly greater on soft gels than on TCPS (Figure 5). On infinitely long channels, the fractional shortening was significantly greater on 10kPa gels than on 35kPa gels (Figure 5). Moreover, fractional shortening of cells on channels in the 35kPa condition was statistically indistinguishable from cells on TCPS. However, measurements of the beating frequency, ∼42-64 beats per minute, did not indicate any statistically significant differences between NRVMs cultured on the hydrogel materials studied. Although lower than clinical values (∼60% for healthy hearts to ∼25% for diseased hearts (33)), fractional shortening of NRVMs cultured on PEG gels reached peak values of 22% fractional shortening on 10kPa hydrogels, significantly greater than that of 13% on 35kPa gels and 7% on TCPS (34).

**Figure 5.**
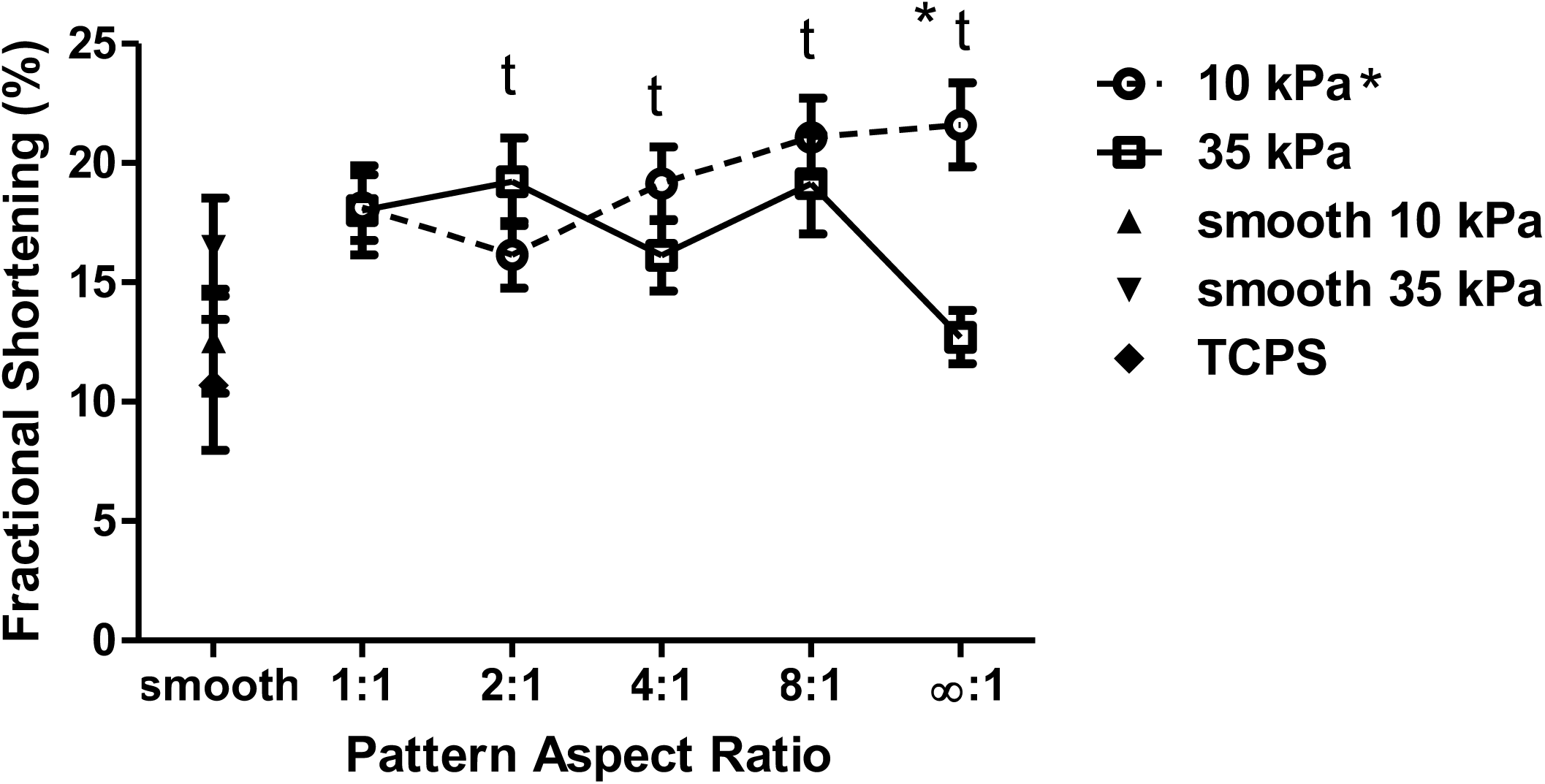
Fractional shortening increases with aspect ratio in softer gels but decreases at highest aspect ratio in stiff gels. Aspect ratios of 2:1 or greater increase fractional shortening relative to TCPS in soft gels. The stiff, channeled pattern significantly reduces fractional shortening relative to soft channels. * next to legend indicates p < 0.05 for regression slope * = p < 0.05 for 10kPa vs. 35kPa, t = p < 0.05 vs. TCPS

### Hydrogel stiffness and patterning attenuate fetal gene expression

Re-expression of fetal genes is a hallmark of cardiac myocyte pathology. Culturing NRVMs on hydrogels significantly reduced the expression of fetal genes, such as *Anf* and *Acta1* (α skeletal actin) compared to TCPS. Relative to TCPS, *Anf* was significantly downregulated on all gel conditions (Figure 6A). *Anf* expression was not different between soft and stiff gels for all pattern aspect ratios studied, but between the 1:1 patterns and smooth gels, there was a ∼2.4-4.4 fold increase in *Anf* expression. *Acta1* expression was also significantly lower on all gel conditions relative to TCPS. Further, *Acta1* expression was significantly lower on soft compared to stiff gels at pattern aspect ratios greater than 2:1 (Figure 6A). However, positive regression slopes for *Anf* and *Acta1* were significant for both soft and stiff hydrogels, indicating increasing expression of these pathologic markers with increasing aspect ratios (Table 1).

**Figure 6.**
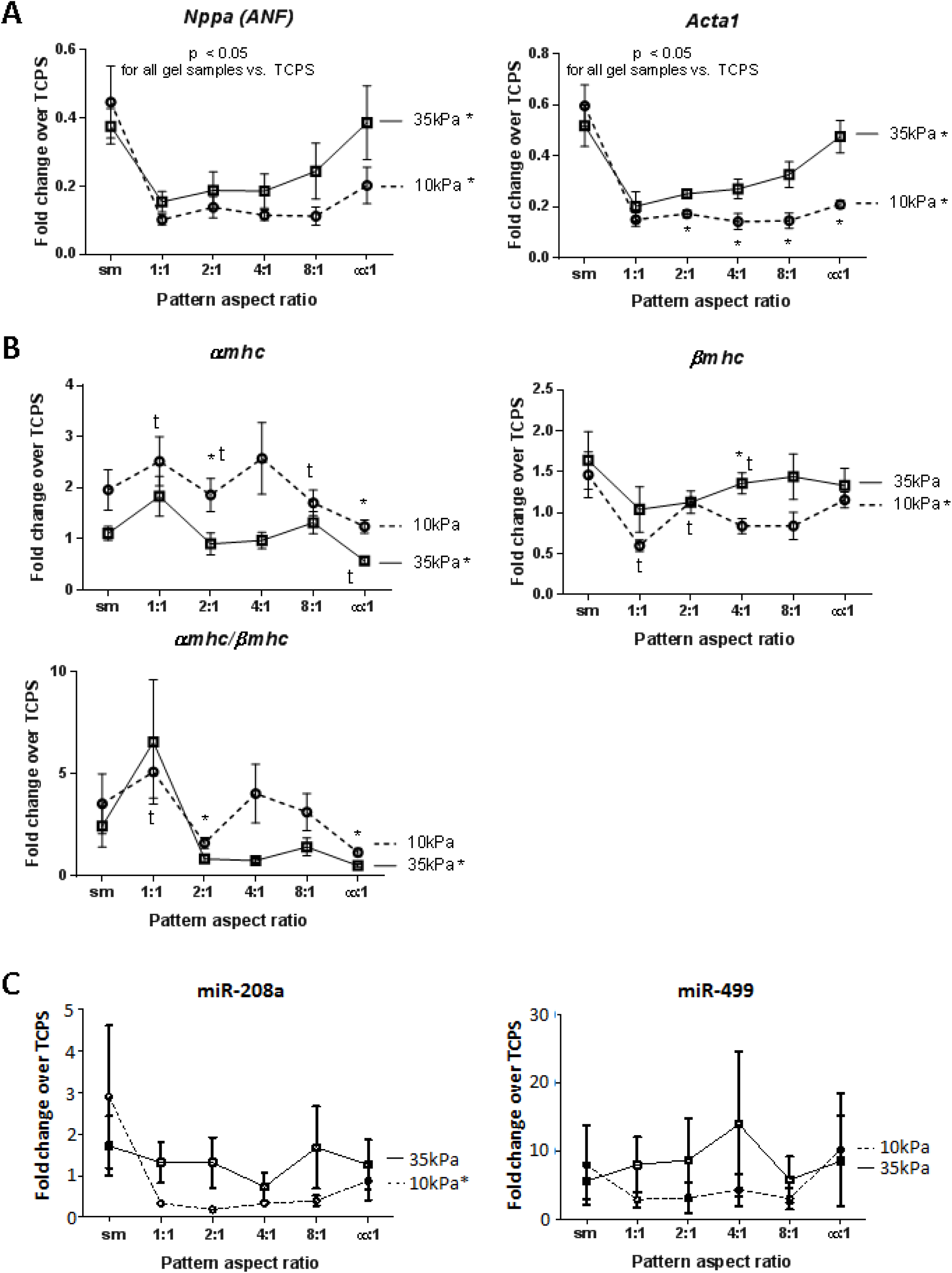
Several pathological fetal genes are downregulated on gels, while pattern aspect ratio significantly affects expression of fetal genes. **A.** Fetal gene expression. *Anf* (*Nppa*) expression significantly increases with pattern aspect ratio as well as a trend for increasing *Anf* expression on stiff, 35kPa gels, vs 10kPa gels. Alpha-skeletal actin (*Acta1*) expression was significantly greater on stiff gels vs soft at aspect ratios greater than 1:1 and increases with increasing pattern aspect ratio. **B.** Alpha-myosin heavy chain (*Myh6*) expression decreases while beta-myosin heavy chain increases on stiff vs soft gels and with increasing aspect ratio. **C.** miR-208a expression decreases on patterned 10kPa hydrogels. miR-208a expression also increases with increasing pattern aspect ratio on 10kPa gels. The fold change magnitude of miR-499 expression on hydrogels vs. TCPS was large but statistically insignificant due to biological variability. * next to legend indicates p < 0.05 for regression slope * = p < 0.05 for 10kPa vs. 35kPa, t = p < 0.05 vs. TCPS

For αMHC, which is generally accepted as a beneficial gene in rodent cardiac myocytes, the regression slope was negative and was significant only for stiff gels. Conversely, the regression slope for βMHC (increases in which are considered pathologic), was positive and was significant only for soft gels. Overall, there was a negative trend between αMHC expression and pattern aspect ratio and a positive trend between βMHC expression and pattern aspect ratio (Figure 6B). Regression slopes were significant for αMHC on 35kPa gels, βMHC on 10kPa gels, and αMHC/βMHC ratio on 35kPa gels. In total, these results suggest that the effect of pattern on myosin heavy chain isoforms is dependent on substrate stiffness.

### Pathological miRNA expression is decreased on patterned 10kPa hydrogels

Given that miR-208a and miR-499 expression is induced in several cardiac disease models and that they may represent potential targets for drug therapies (35–37), we analyzed their expression as a function of NRVM culture conditions (Figure 6C). Significant observations included a reduction in NRVM miR-208a expression when cultured on patterned (1:1 through 8:1), 10kPa hydrogels compared to TCPS; these results suggest a protective effect from a combination of substrate patterning and decreased substrate stiffness. However, the regression analysis indicated a positive slope for miR-208a expression with increasing aspect ratio on soft gels (Figure 6C). There was also a nonsignificant, but large magnitude trend for higher miR-499 expression on 35kPa hydrogels. These trends are consistent with the gene expression patterns observed in Figure 6A and B and together suggest that patterned hydrogel substrates could serve as platforms to model cardiac disease processes at multiple biological levels.

### Hydrogel patterning and stiffness do not significantly affect calcium handling genes

When simultaneously modifying substrate stiffness and patterning, expression of the calcium handling genes Cav1.2 (*Cacna1c*) and Serca2a (*Atp2a2*) was not significantly different between experimental groups; however, there was a non-significant trend for higher *Cav1.2* and *Serca2a* expression on 10kPa gels (Figure S1 and Table 1). During aging and disease, *ATP2a2* is downregulated (38), while *Cav1.2* expression is reduced in pressure overload conditions (39). Varying substrate stiffness between 10kPa and 35kPa while presenting a range of pattern aspect ratios did not replicate the fold changes observed in disease models. The results suggest that a larger range of substrate stiffness or pattern shapes may be necessary to produce changes in calcium handling genes that are observed in disease.

### Matrix remodeling genes are differentially regulated by culture on hydrogel substrates

Markers of ECM remodeling, such as connective tissue growth factor (*Ctgf*) and collagen type 1 alpha 1 (*Col1a1*) expression, were differentially regulated by substrate stiffness and by pattern aspect ratio (Figure S2). *Ctgf* expression on all gel samples was significantly reduced by at least two-fold relative to TCPS. *Ctgf* expression was also significantly higher on stiff gels for several pattern geometries, and regression slopes were significant for both soft and stiff gels samples. These results suggest that both substrate stiffness and pattern aspect ratio affect *Ctgf* expression. Neither pattern aspect ratio nor gel stiffness significantly affected matrix metalloproteinase 2 (*Mmp2*) expression; however, *Mmp2* expression on soft gels with 1:1 and 2:1 patterns was significantly greater than on TCPS (Figure S2). *Col1a1* expression was significantly lower on soft gels than on TCPS, but differences between soft and stiff gels were not significant. Regression slopes for *Col1a1* expression were also not significant for soft and stiff gels.

## Discussion

*In vivo*, cardiac myocytes experience both mechanical and topographical changes to their microenvironment during development, physiological remodeling, and disease (10, 12). Although many diverse animal models exist to test hypotheses generated from clinical observations, some hypotheses require *in vitro* systems as they offer much higher levels of control than an animal. However, *in vitro* models have lagged behind animal models in terms of providing a flexible system that can model both physiologic and pathologic cardiac states. Thus, a critical need in the field of cardiac biology is an *in vitro* system that recapitulates the (1) physical, (2) morphological, (3) functional, and (4) molecular features of both healthy and diseased microenvironments.

NRVMs are the most commonly used cardiomyocyte for *in vitro* studies, and several studies demonstrate that NRVMs adapt to changes in substrate stiffness or patterns by regulating their gene expression, morphology and contractility (15, 40). However, the vast majority of these studies investigate the effect of modifying one particular aspect of the cellular microenvironment amongst a multitude of factors that can change during normal development and disease processes. In this work, we developed a photodegradable and photopatternable hydrogel platform with varying micropatterns and stiffnesses (schematized in Figure 1) to evaluate whether this system could faithfully model both physiologic and pathologic states by comprehensively examining all four stated aspects of cardiomyocyte biology. In addition, the feature sizes were micron size and smaller than a typical NRVM. Thus, rather than physically constraining the NRVM artificially with topography, the hydrogels promote cell-interactions and allow the NRVMs to spontaneously align based on their underlying focal adhesions with the substrate. In this way, the hydrogel recapitulates aspects of an organized ECM-interaction and its influence on mechanotransduction. We hypothesized that the conditions in which the cardiomyocytes were best aligned on the softer gels (10kPa) would preserve the most physiologic phenotype because the healthy neonatal myocardium is characterized by this modulus and by well-aligned cells.

Initial experiments revealed that NRVMs cultured on patterned hydrogel substrates formed mature sarcomeres. Spontaneous contraction was also observed in all experimental groups. Importantly, cell density appeared similar across all groups. These observations suggested to us that NRVMs cultured on PEG hydrogels maintained a cardiac phenotype and that our hydrogel system was suitable for more in-depth characterization of cells cultured using this platform. Biochemical and image analysis measurements indicated that both substrate stiffness and topography are significant factors that ultimately determine alignment, morphology and gene expression of NRVMs (Table 1). For many of the readouts that we investigated, the largest fold changes were observed only when varying both substrate stiffness and patterning. For example, although cellular alignment was highest on 35kPa gels with ∞:1 aspect ratio patterns, fractional shortening was actually the lowest under those conditions. *Anf* and *Acta1* expression was lowest on 10kPa gels, but increased with increasing aspect ratio on gels regardless of the underlying stiffness. These results suggest that optimal levels of gene expression and contractility may occur at an intermediate pattern/aspect ratio on a soft hydrogel and that incorporating multiple microenvironment cues may increase the response range of NRVMs and lead to fold changes in assay results at clinically relevant levels. Importantly, this work emphasizes the fact that all aspects of cardiomyocyte health must be assessed, as relying on characteristic morphological changes alone can be misleading.

### (1) Physical features: intercellular alignment and substrate modulus

Myocyte disarray is a key feature of cardiac disease and cells plated on traditional, flat surfaces are disarrayed in a manner similar to that observed in animal and human disease (1, 41, 42). Prior studies have also found that eroding topographical patterns affects the alignment of NRVMs on smooth surfaces and surfaces with only one or two different pattern dimensions (21, 22, 43). In contrast, examining a gradient of pattern aspect ratios provides the ability to mimic intermediate conditions, rather than only extremes. In addition to myocyte disarray, the stiffness of diseased hearts also increases. Using photodegradable PEG hydrogel substrates, we generated a range of pattern aspect ratios on both soft (10kPa) and stiff (35kPa) surfaces. The ability to simultaneously deliver multiple microenvironmental cues, namely stiffness and patterning, greatly expands the possible experimental space and provides the ability to examine how multiple features of disease interact. Few studies have varied both substrate stiffness and patterning. Notably, McCain *et al.* used microcontact printing to generate large, NRVM-sized fibronectin islands of varying aspect ratios on soft and stiff polyacrylamide hydrogels in order to generate cells of varying aspect ratios. They observed that shorter aspect ratio cells generated the most systolic work on 90kPa gels while longer aspect ratio cells generated the most systolic work on 13kPa gels (40). Extending from this work, Ribeiro et al. cultured human cardiomyocytes differentiated from induced pluripotent stem cells (iPSCs) on 10kPa polyacrylamide hydrogels patterned with 2000 μm^2^ matrigel adhesion islands to constrain cells to different aspect ratios. They demonstrated that such a platform resulted in better differentiation and higher mechanical output (29).

In contrast, our study evaluated whether cardiomyocytes would spontaneously adapt to micropatterned substrates. We photo-eroded small micropatterns (∼1000µm^2^), less than the size of single cells, to direct cell alignment without forcing a particular cell shape and aspect ratio through adhesion islands, as is the common approach in published studies (29, 40). We interpret the spontaneous molecular, structural, and functional changes we observed to represent a true signal-response relationship between the environment and the myocyte that may more accurately reflect *in vivo* cardiac remodeling. For example, fold changes in percent alignment observed between patterned and smooth substrates were consistent with changes in myocyte alignment in studies of wildtype mice (∼46% aligned) and mice with hypertrophic cardiomyopathy (∼25% aligned) (32). Myocyte disarray was modeled using a range of micropattern aspect ratios, and NRVMs cultured on soft and stiff hydrogels exhibited similar changes in disarray as observed in *in vivo* systems (Figure 3). Our results indicate a greater percentage of cells aligned on the 35kPa ∞:1 substrate than the 10kPa ∞:1 substrate. This finding may seem to contradict our initial assumption that the 10kPa substrate modeled a healthier condition. However, we also observed highest function on the 10kPa ∞:1 substrate. Disease progression is a complicated multifactorial process where all cells start at the same state and cells in the diseased tissue remodel and respond to their changing conditions. In a comparably simple *in vitro* system, the “health” range of cell alignment may be different. Nonetheless, our patterned substrates could induce cell alignment significantly greater than that of smooth surfaces.

### (2) Morphological features: cellular aspect ratio

During heart failure, the morphology of human cardiac myocytes changes as the cells elongate, and aspect ratios have been known to increase by a factor of 1.8 (34, 44, 45). Moreover, increased aspect ratios can result from long-term aerobic training while decreased aspect ratios can result from resistance training. Therefore, aspect ratio should not be viewed as a singular metric of cardiomyocyte health. We took a multifactorial approach and modified substrate stiffness and topographic pattern aspect ratios concomitantly. This resulted in cell aspect ratios that spanned a 2.2 fold range, covering fold changes observed in human samples and rodent models. It is also interesting to note that the NRVM aspect ratios on 10kPa hydrogels alone appeared to cover a slightly broader dynamic range than on stiff hydrogels. Simultaneously varying substrate stiffness and patterning resulted in significantly greater F-actin alignment on stiff hydrogels than on TCPS (Figure 4). These observations suggested to us that in addition to the significant effect of the pattern’s aspect ratio on F-actin alignment, the effect of substrate stiffness may depend on the underlying pattern. While the bending stiffness and available binding area of patterns may also influence cell morphology and F-actin organization, systematically modifying these factors, such as using more than two stiffnesses, was beyond the scope of the current study and is the subject of future experiments. Although neonatal cells were used, simultaneous modulation of substrate stiffness and pattern aspect ratio produced a range of cell aspect ratios that span values measured in both healthy adult individuals and patients with ischemic cardiomyopathy (45).

### (3) Functional features: fractional shortening

Fractional shortening values that were measured while varying substrate stiffness and patterning ranged from ∼13% to ∼22% for a dynamic range of 1.7 fold. These values are similar to those reported elsewhere for NRVMs cultured on 8kPa polyacrylamide hydrogels (46). Clinical fractional shortening values based on *in situ* measurements of ventricular geometry range from up to ∼60% for healthy hearts to ∼25% for diseased hearts (33); however, isolated cells in culture often exhibit much lower values of ∼7% (34). On 10kPa hydrogels, fractional shortening increased with pattern aspect ratio, while on 35kPa hydrogels there was a trend for decreasing fractional shortening with increasing pattern aspect ratio. Fractional shortening dropped precipitously between 8:1 and channel features on 35 kPa gels. It is worth noting that the cells need to form focal adhesions with the matrix, and these would be expected to be sparser in the short dimension on channels where they are less likely to encounter the substrate. This, coupled with the pathologic changes in gene expression (particularly the decreased αMHC and increased βMHC expression) could account for this decrease in function. These results suggest that substrate stiffness significantly affects fractional shortening when cells are aligned with high aspect ratio patterns. Another observation is that fractional shortening was greatest at an intermediate value of effective stiffness, a function of substrate stiffness and patterning. Other studies have shown optimal contractility on smooth substrates with an intermediate stiffness, ∼10kPa (16, 47). Simultaneous delivery of multiple microenvironmental cues revealed potential interacting variables such as available cell growth area, effective substrate stiffness, and pattern geometry. These experimental parameters represent opportunities to better design optimal cell culture platforms. Investigating how these factors affect NRVM contractility is a subject of ongoing studies.

### (4) Molecular features: gene expression signatures

Fetal genes are expressed during development but are downregulated during normal postnatal growth and adulthood (48). *Anf* and *Acta1* are upregulated during cardiac disease and the expression of both was attenuated in NRVMs cultured on all PEG hydrogels relative to TCPS. In comparing 1:1 patterned to smooth hydrogels, *Anf* expression increased ∼2.4-4.4 fold, thus mimicking fold changes seen clinically when volumetric overload of the heart causes stretching of the myocardium and increased matrix stiffness sensed by the myocytes (49). Expression of *Acta1* has been positively correlated with contractile function (50), a phenomenon we observed with increasing aspect ratio on 10kPa substrates. However, our study also supports the observation that *Acta1* expression increases with increasing disease burden as we measured the highest levels of *Acta1* on 35kPa gels with ∞:1 patterns, where we also measured lowest fractional shortening (Figure 5). Notably, we measured as great an increase in *Anf* and *Acta1* expression due to increasing the pattern aspect ratio as due to increasing the substrate stiffness. These data suggest that patterning had a greater effect on *Anf* and *Acta1* when presented in stiff environments compared to soft.

Matrix remodeling, alterations in normal calcium handling, and a decrease in the αMHC/βMHC ratio are also seen in cardiac disease (51). Connective tissue growth factor (CTGF) is induced during heart failure (52). It is upstream of many ECM remodeling pathways and the accumulation of matrix proteins, such as collagens, results in fibrosis further leading to dysfunction and even sudden cardiac death. Increasing pattern aspect ratio led to increasing *Ctgf* expression regardless of substrate stiffness (Figure S2). However, the expression of *Col1a1* was significantly lower than TCPS only on soft hydrogels. These findings suggest that global ECM remodeling is sensitive to small changes in substrate stiffness. Although we did not observe differential expression of the calcium handling genes *Cav1.2* and *Serca2a* in any of our conditions (Figure S1), a larger range of substrate stiffness may be required to observe the same changes found in other studies. αMHC expression was greater on soft, 10kPa gels than on stiff gels. While we measured generally higher (and thus more physiologic) αMHC/βMHC ratios on soft gels (Figure 6), the αMHC/βMHC ratio appears to decline beyond the 4:1 aspect ratio and ultimately approach that of the 35kPa gels. This suggests increasing pathologic state with increasing aspect ratio. We also found significantly decreased expression of miR-208a on 10kPa gels relative to TCPS but a positive regression slope with increasing aspect ratio. Increased human patient serum levels of miR-208a and miR-499 have been observed after myocardial infarction (53), and transgenic overexpression of each is sufficient to cause cardiac hypertrophy and dysfunction in mice (37, 54). Once again, high aspect ratios begin to counteract the beneficial effects of soft substrates. These results suggest that softer substrates promote greater contractility as well as reduced pathological gene expression at intermediate aspect ratios, as summarized in Figure 7.

**Figure 7.**
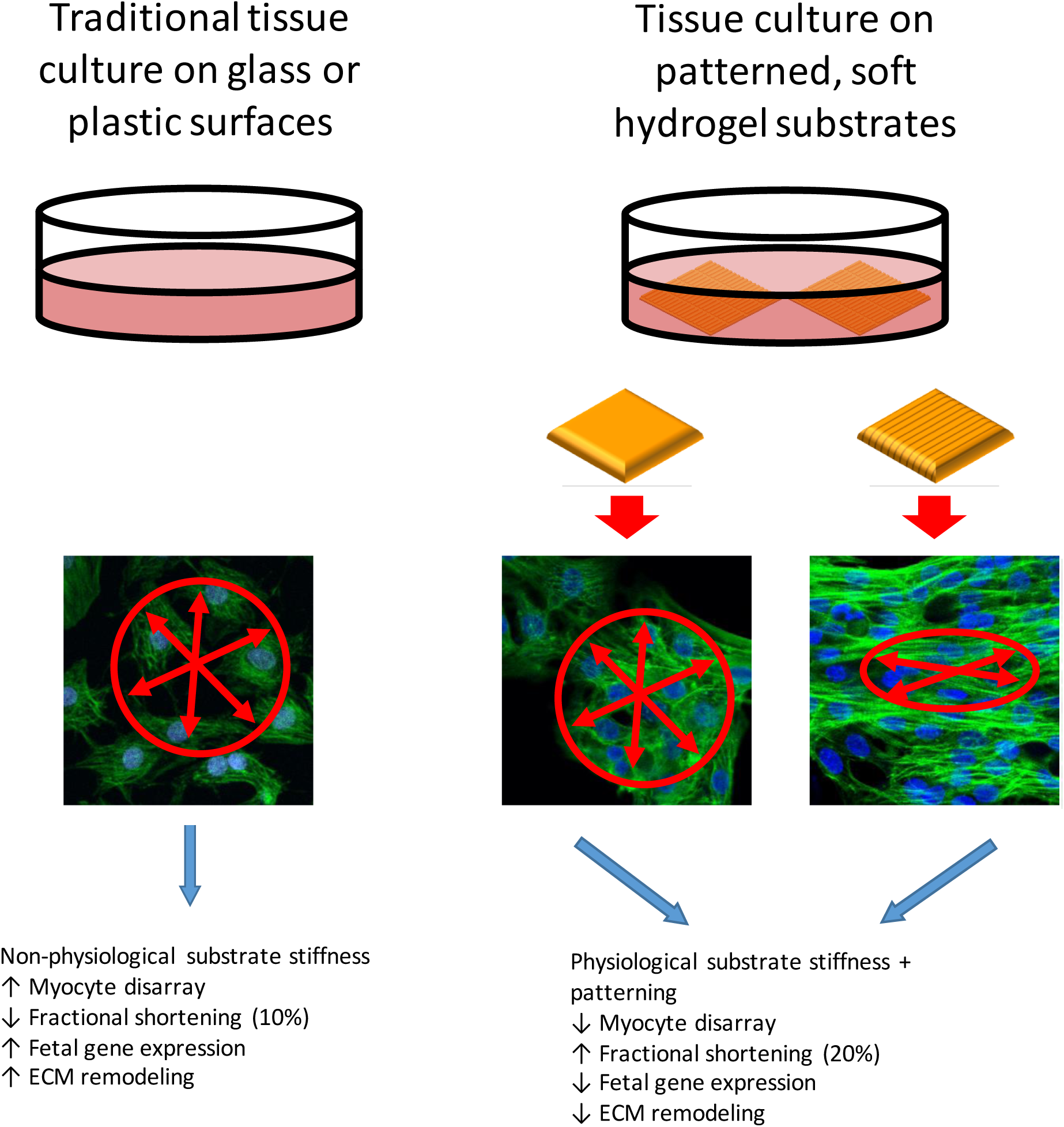
Substrate stiffness and topographical cues significantly affect NRVM physiology. Culturing NRVMs on 10kPa patterned hydrogels resulted in attenuated pathological gene expression as well as increased fractional shortening. Using a range of patterns and substrate stiffnesses, we were able to capture both physiological and pathological states.

#### Study Limitations and Future Directions

This work along with the previous studies referenced herein indicate that substrate modulus and topography are strong stimuli for cardiomyocyte adaptation to environmental changes and that the culture platform can be manipulated to mimic both pathologic and physiologic cardiac states. The ultimate goal in modeling cardiomyocyte biology, however, is to establish a system conducive to culturing adult human cardiomyocytes. The study presented here uses NRVMs, which are both a different species and a different developmental stage. Unfortunately, current iPSC-derived human cardiomyocytes more closely resemble fetal cardiomyocytes and often these cultures differentiate to mixtures of atrial, ventricular, and nodal cardiomyocytes (55). Ribeiro and colleagues demonstrated that gel-based culture platforms with patterned adhesion islands could improve differentiation of iPSC-derived human cardiomyocytes, although their system imposes cellular morphology through adhesion islands with no resulting gene expression differences between patterned and unpatterned substrates (29). Given that our system simply provides topographical cues with significant consequences to multiple aspects of cellular mechanics and physiology, it would be interesting to evaluate whether further improvements to iPSC differentiation, particularly with respect to cell fate, could be achieved with our system. Another key consideration is the three dimensional environment of the culture platform. In our current system as well as many published systems, cells are cultured on top of gels. However, the heart is a three-dimensional organ and ventricular myocytes are typically encased in the extracellular matrix. In a two-dimensional culture platform, only one face of the cell is in contact with the substrate, creating a differential with respect to mechanical and chemical cues. Thus a logical extension is to move towards a three-dimensional hydrogel-based culture system, an active area of investigation. Indeed, Ronaldson-Bouchard and colleagues have achieved vast improvements in maturation (ultrastructure, gene expression, contractile properties, metabolic profiles, and calcium handling) of *in vitro* differentiated hiPSC-derived cardiomyocytes by combining three-dimensional fibrin hydrogel platforms with dynamic electrical stimulation regimes beginning early in differentiation (56). Combining this approach with a tunable hydrogel system whose stiffness could be matured from a more pliant, fetal range to a stiffer, more adult-like range while the cells differentiate is intriguing. In addition, translating 2D topographical patterning to 3D cell culture environments is an area of growing interest, and photopatternable materials, as the one used here, provide specific advantages over micromolding or microprinting techniques when combined with single and two photon laser lithography. Such a system would also be amenable to modeling interfaces, such as between the border zone and the infarct zone of an infarcted heart.

We measured contractility as an indicator of cardiomyocyte function, but another key functional measurement is the traction forces the cell exerts on the substrate as it contracts. While this technique has been well-refined for cells on planar surfaces, it is still being developed for cells on three-dimensional substrates (57). Non-planar substrates lead to large error in the z direction with variable point spread functions, making reliable analysis with typical equipment difficult. Advances in hardware and analytical approaches should make this a more tractable problem in the future.

#### Conclusions

Cardiac myocytes sense and interact with their microenvironment through a wide range of biochemical and morphological responses (Figure 7). Here, biochemical and biophysical approaches were used to investigate how NRVMs respond to multiple, concurrent microenvironmental cues, namely substrate stiffness and patterning. In contrast to previous studies that examined the response of NRVMs to only a single factor, this study demonstrated that both substrate stiffness and topographical patterning synergistically affect cellular morphology, NRVM gene expression profiles and contractility. In many cases, we observed the effects due to substrate stiffness only at certain pattern aspect ratios. This study underscores how cells can integrate multiple microenvironmental cues, and that concurrent signals can synergistically alter cell function. Because of the complexity of the cellular response to matrix signals, it is essential to test several experimental factors in *in vitro* disease models, since results from modifying only one factor may mask the effects of other critical factors. In addition, many studies evaluating the cellular response to environment measure few outputs, such as cell morphology. However, this study reveals that a broad range of cellular metrics (i.e., physical, morphological, functional, and molecular) must be tested in order to fully evaluate the health of a cardiomyocyte: gross readouts such as cell morphology can be misleading when considered alone. Importantly, simultaneous presentation of substrate stiffness and topographical cues replicates fold changes in cell morphology and gene expression observed *in vitro* as well as in animal models of cardiac disease. The collective results suggest that differences in the microenvironmental stiffness and patterns that influence cell-matrix interactions are both important in regulating cardiac myocyte gene expression and contractile response. Recreating multiple aspects of *in vivo* systems using bioscaffolds with tunable properties for *in vitro* experiments provides complementary information that allows investigators to more comprehensively understand how cells respond to physiological and pathological microenvironments.

## Materials and Methods

### Hydrogel formulation

A photodegradable PEG crosslinker (PEGdiPDA) was synthesized as previously described (58). PEGdiPDA (M_n_ ∼ 4070 g/mol, 8.2 wt%) was copolymerized with poly(ethylene glycol) monoacrylate (M_n_ ∼ 400 g/mol, 6.8 wt%, Monomer-Polymer and Dajac Laboratories, Inc) via a radical initiated chain polymerization. Gelatin (100 bloom, 1mg/mL, MP Biomedicals) was included in the monomer solution to promote cell-matrix interactions and adhesion. Gelation was initiated through a redox reaction by combining 0.2M ammonium persulfate with 0.1M tetraethylmethylenediamine. The monomer solution was briefly vortexed and pipetted between a glass slide and glass coverslip that was functionalized with acrylate groups (59) to covalently link the final hydrogel to the coverslip. After gelation (∼5 minutes), hydrogels were briefly immersed in phosphate buffered saline (PBS), separated from the glass slide, and stored in PBS at 4°C until use. Rheological measurements of the initial, stiff (35kPa, Young’s modulus) gels were similar to previous studies from our group (60).

### Generation of micropatterned substrates

A custom photomask was used to control the pattern of light illumination and generate topographical patterns on polymerized gels. The originally fabricated stiff gels (35kPa) were placed in contact with the selected photomask and subsequently irradiated with collimated 365nm light (Omnicure) at 15mW/cm^2^ for 300s. The photomask allowed transfer of micropatterned rectangles to the gel substrate that were 5µm apart with geometries of 5 µm × 5 µm, 10 µm × 5 µm, 20 µm × 5 µm, 40 µm × 5 µm and 5 µm wide channels corresponding to aspect ratios of 1:1, 2:1, 4:1, 8:1 and infinity:1 (∞:1) (Figure 1). To generate soft patterned substrates, a set of patterned gels was then uniformly irradiated, without a photomask, for 300s. This additional exposure step generated substrates with a reduced crosslinking density, corresponding to a 10kPa gel stiffness. Smooth, unpatterned, 35kPa and 10kPa gels were also used. The irradiation time used to generate soft surfaces, via controlled photodegradation of the hydrogel crosslinks, was similar to the time scale and values reported in previous studies (60). In previous studies from our group, we validated rheometry measurements of irradiated gels by atomic force microscopy and validated pattern uniformity by two-photon confocal laser scanning microscopy (61).

### Cell isolation and culture

Neonatal rat ventricular myocytes were isolated according to previously published procedures (62). Unless noted, all reagents were purchased from Sigma. Briefly, hearts were excised from 1-3 day old Sprague-Dawley rat pups. The atria were removed from the hearts and discarded. The remaining ventricle sections were minced using scissors and digested in trypsin. After pre-plating for 2 hours, NRVMs in suspension were collected and counted. Primary cell isolates were seeded at ∼50,000 cells/cm^2^ on gel samples or on gelatin coated standard tissue culture polystyrene (TCPS) plates. During the first 24 hours of culture, cells were plated in growth media containing Minimum Essential Medium (MEM) with Hank’s salts (Gibco), 5 vol% calf serum, 50U/mL penicillin, 2µg/mL Vitamin B-12, and 30nM bromodeoxyuridine (BrdU). After 24 hours, cells were washed, and the media was changed to one consisting of MEM, 10µg/mL insulin, 10µg/mL transferrin, 0.1 wt/vol% bovine serum albumen (BSA), 50U/mL penicillin, 2µg/mL Vitamin B-12, 30nM BruU, and 10 vol% fetal bovine serum (FBS). For subsequent experiments, cells were collected or stained at day 4. All animal procedures were approved by the Institutional Animal Care and Use Committee at the University of Colorado.

### Fractional shortening

Videos of spontaneously contracting cardiac myocytes were collected through a 40× objective (1.0 NA) at 50 frames per second using a high-speed camera (HiSpec1 G2, Fastec Imaging) on an upright widefield fluorescence microscope (Examiner.Z1, Zeiss). While imaging, cells were maintained in warm Tyrode’s solution, and before imaging, cells were pre-incubated with 2µM CellTracker Red CMPTX (Life Technologies) for visualization under fluorescence and for later segmentation and contractility measurements.

### Cell staining

Cells were fixed in 4% paraformaldehyde (Electron Microscopy Sciences) for 5 minutes, washed in PBS and permeabilized in 0.1% Triton X-100 (Fisher) for 3 minutes. F-actin and nuclei were stained with phalloidin conjugated to an Alexa Fluor 488 fluorophore (Life Technologies) and 4’,6-diamidino-2-phenylindole, DAPI (Life Technologies), according to manufacturer’s instructions. All samples were imaged on a confocal microscope (Zeiss LSM 710 NLO) and processed using ImageJ (National Institutes of Health, USA).

### Image analysis

Organization of F-actin fibers was quantified with Matlab (MathWorks, Natick, MA) to calculate the percent of fibers aligned with the mean fiber angle (63, 64). Images of F-actin fibers were thresholded using Otsu’s method, windowed by applying a 2D Tukey window, and a fast Fourier transform (FFT) was performed. The power spectrum of the FFT was used to generate a histogram of frequency intensities between -89° and 90°. Aligned F-actin fibers were defined as those whose orientations were ±20° of the mean fiber angle. Thus, the working range of percent alignment was at least ∼22% for randomly oriented fibers and 100% for samples where all fibers were within ±20° of the mean angle. The percent alignment of gel samples was further normalized to the mean percent alignment of cells cultured on TCPS.

Percent fractional shortening (FS) was calculated from videos of contracting cells according to

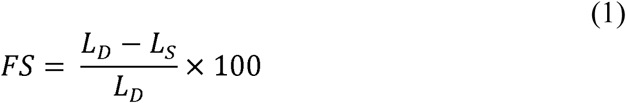

Here, *L*_*S*_ is the major axis of a maximally contracted cell and *L*_*D*_ is the major axis of a fully relaxed cell. Videos of single contracting cells were analyzed using a custom Matlab script. Individual frames from each video were thresholded and individual cells were identified. The area was calculated using a built-in Matlab function, regionprops. Because individual, non-confluent cells were analyzed, the outline of the cell was easily segmented and outlined. The boundary of the cell was fit to an ellipse using built-in Matlab functions. The cell aspect ratio was calculated as the ratio of the major axis length to the minor axis length.

The degree of cell alignment with respect to the pattern orientation was measured using a custom ImageJ macro. The alignment of the cell was assumed to follow the alignment of the nucleus (65). The major axis of the pattern was manually measured and followed by an automated measurement of the nuclear orientation from the DAPI channel image. The alignment of cells with respect to the pattern orientation was calculated by subtracting the measured angle of cell nuclei from the manually measured major axis of the pattern. Mean angle differences were calculated using circular statistics (66).

### Quantitative polymerase chain reaction (qPCR)

RNA was isolated using a commercial kit (RNAmicro kit, Zymo Research), according to the manufacturer’s instructions. RNA concentration and quality were measured using a spectrophotometer (Nanodrop 1000, Thermo Scientific), and samples were stored at -70°C until further use. For mRNA analysis, cDNA was synthesized using a commercial kit (SuperScript III First-strand synthesis kit, Invitrogen) using random hexamers according to kit instructions. For qPCR, a 10µL reaction volume mastermix was prepared, containing forward and reverse primers (420nM each) and SYBR green (Applied Biosystems). The mastermix was loaded onto 96 well plates and reactions took place on a thermal cycler (Bio-Rad) programmed according to SYBR green instructions. Ribosomal protein 30 (RPL30) was used as the reference gene and relative quantification with the Pfaffl method was used to determine gene expression changes. Target genes are listed in Table S1.

For miRNA analysis, Taqman®-based assays were performed according to the manufacturer’s instructions (Applied Biosystems). Briefly, 6.67 ng total RNA was reverse transcribed in 10μL using a TaqMan® MicroRNA Reverse Transcription Kit (Applied Biosystems catalog #4366596). miRNA expression was assessed with 20μL reactions prepared from the TaqMan® Universal PCR Master Mix, no AmpErase® UNG kit (Applied Biosystems catalog #4324018) and the appropriate TaqMan MicroRNA assay (catalog #4427975) on a Bio-Rad thermal cycler with cycling conditions according to the TaqMan instructions. Relative quantification based on the ΔΔC_T_ method was performed with U6 RNA as the reference gene.

### Data analysis

Each experiment consisted of at least 9 replicates from several independent cell isolations. Data are presented as mean ± standard error of the mean (SEM). The effects of substrate stiffness and pattern geometry were analyzed by defining pattern geometry as an ordered factor and applying general linear models in R (67). Analysis of variance (ANOVA) was used to estimate the effect of pattern geometry and substrate stiffness on the experimental outputs. Regressions were used to separately estimate the effect of pattern geometry on cells that were cultured on either soft or stiff substrates. Pattern aspect ratios were entered as ordered categorical factors in the following order: smooth, 1:1, 2:1, 4:1, 8:1 and infinity:1 (∞:1). Slopes that were significantly greater than zero indicated a significant effect in the cellular response with increasing aspect ratio in the topography of the hydrogel surface. *Post-hoc*, pairwise comparisons were also performed, and p-values adjusted using Bonferroni’s correction. Cell alignment data were analyzed using circular statistics, and the equal kappa test was used to determine whether standard deviations were statistically different from smooth and patterned surfaces. While biological systems are often nonlinear, general linear models provide conservative measures of experimental effects and minimize chances of overfitting the data. Effects, slopes, and comparisons were considered significant at p < 0.05.

## Acknowledgements

We thank Ann Robinson for assistance with NRVM cell isolation, and Ciera N. Dolechek for technical assistance. We also thank Dr. Chelsea Magin for insightful discussion.

## Sources of funding

This work was supported by the National Institutes of Health (GM029090), the Howard Hughes Medical Institute, the National Science Foundation Postdoctoral Research Fellowship in Biology under Grant No. (1307559), the American Heart Association Postdoctoral fellowship to W.W. (13POST14730015), and the American Heart Association Undergraduate Student Research Program to E.S.C. (16UFEL31660000).

## Disclosures

None

