## Supplementary figures and images for "Cardiac myocytes respond differentially and synergistically to matrix stiffness and topography"

### Supplemental Figures

#### Slide 1
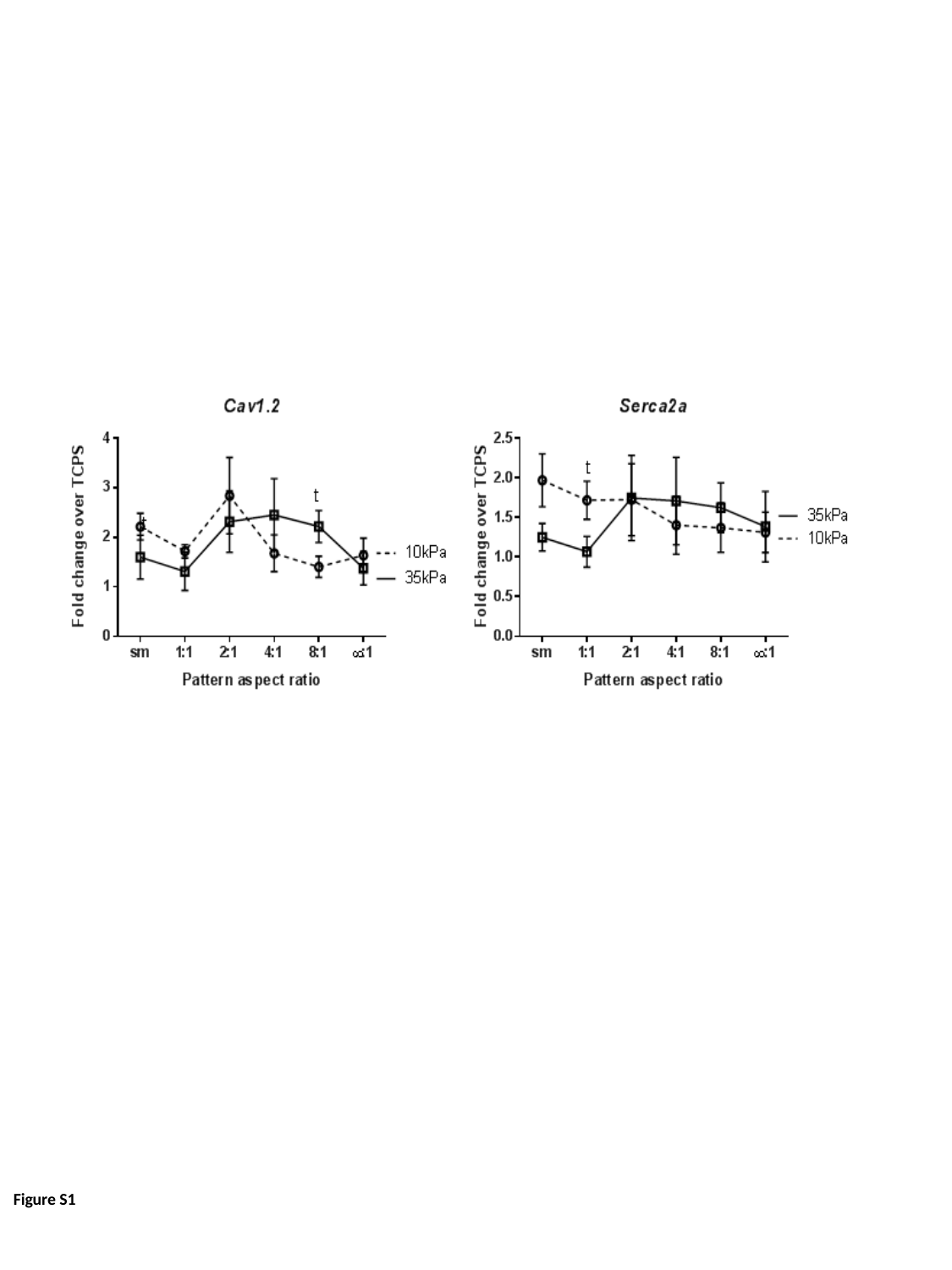

Figure S1

#### Slide 2
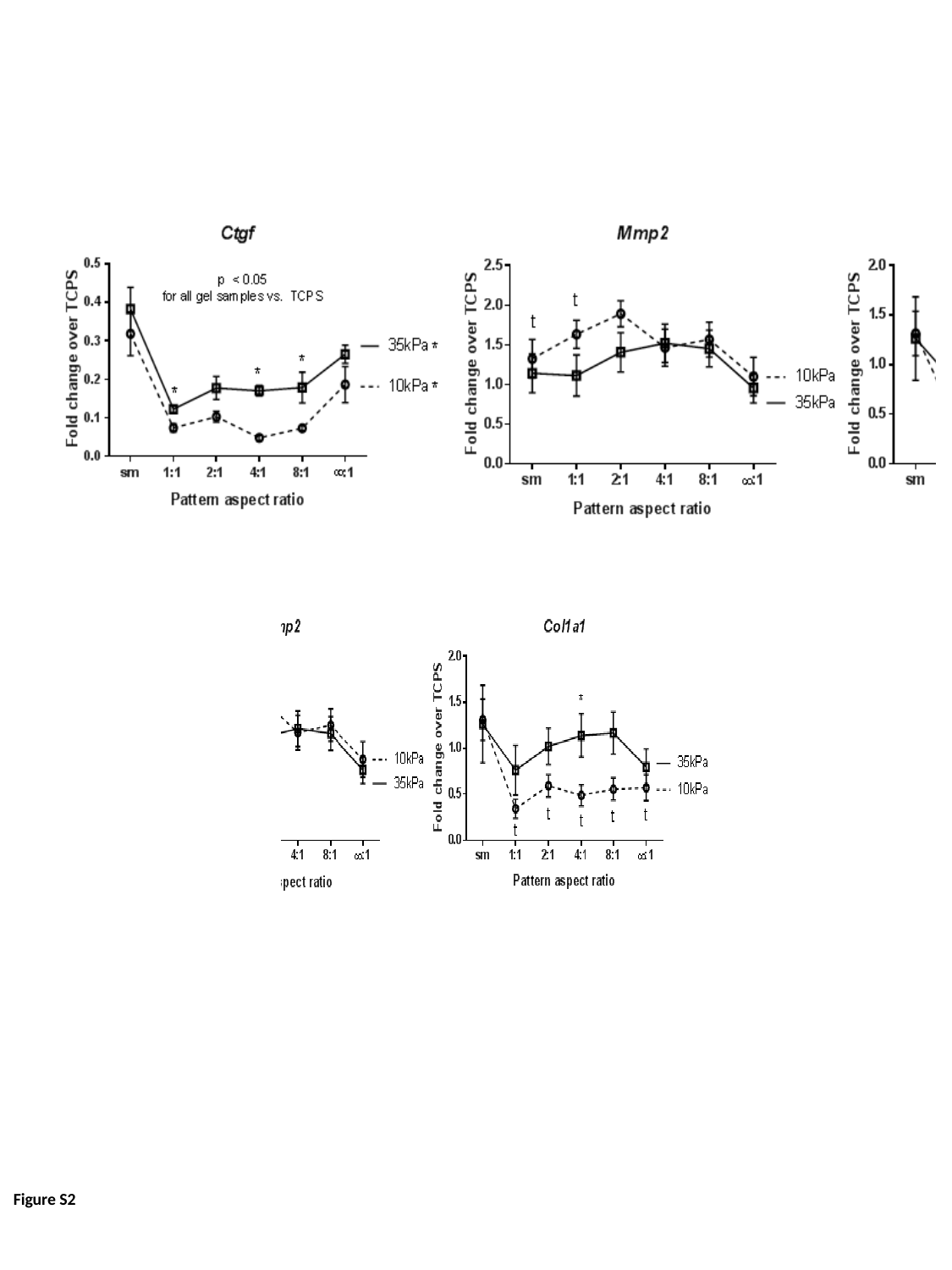

Figure S2
