## Supplemental Figure Legends for "Cardiac myocytes respond differentially and synergistically to matrix stiffness and topography"

**Figure S1: Calcium handling genes did not significantly vary in most experimental groups.** Expression of the calcium handling genes *Cav1.2* and *Serca2a* was not significantly affected by substrate stiffness nor patterning.

**Figure S2: Substrate stiffness and topographical cues affect some ECM remodeling genes.** Pattern aspect ratio and substrate stiffness significantly affects *Ctgf* expression; however, ECM remodeling may be taking place through mechanisms independent of collagen1 and Mmp2 regulation.

* next to legend indicates $p < 0.05$ for regression slope

* = $p < 0.05$ for 10kPa vs. 35kPa, t = $p < 0.05$ vs. TCPS

**TABLE S1:** Gene symbols and primer sequences

| Gene name (abbreviation in paper) | Gene symbol | Forward primer | Reverse primer |
|---|---|---|---|
| Ribosomal protein L30 (*Rpl30*) | *Rpl30* | CTTGGCGTCTGATCTTGGTT | AAGTTGGAGCCGAGAGTTGA |
| Atrial natriuretic factor (*Anf*) | *Nppa* | GCGAAGGTCAAGCTGCTT | CTGGGCTCCAATCCTGTCAAT |
| α-myosin heavy chain (αMHC) | *Myh6* | GCGCCAAGCAGAAAATGCAC | TGTGGGATAGCAACAGCGAG |
| β-myosin heavy chain (βMHC) | *Myh7* | CGCTCAGTCATGGCGGAT | GCCCCAAATGCAGCCAT |
| Calcium channel, voltage-dependent, L type, alpha 1C subunit (*Cav1.2*) | *Cacna1c* | CGATGTGAAGGCACTGAGAG | GATGATGACGAAGAGCACGA |
| ATPase, Ca++ transporting, cardiac muscle, slow twitch 2 (*Serca2a*) | *Atp2a2* | ACCTGGAAGATTCTGCGAAC | AATCCTGGGAGGGTCCAG |
| α-skeletal actin (*Acta1*), | *Acta1* | TGAAGCCTCACTTCCTACCC | CGTCACACATGGTGTCTAGTTTC |
| Matrix metalloproteinase 2 (*Mmp2*) | *Mmp2* | GGGCACCTCTTACAACAGC | AGTGGACATAGCAGTCTCT |
| Collagen type 1 (*Col1a1*) | *Col1a1* | GACTGTCCCAACCCCCAAAA | ACTTCTGCGTCTGGTGATACATA |
| Connective tissue growth factor (*Ctgf*) | *Ctgf* | AGCAGCTGGGAGAACTGTGC | ACTGCTTTGGAAGGACTCGC |

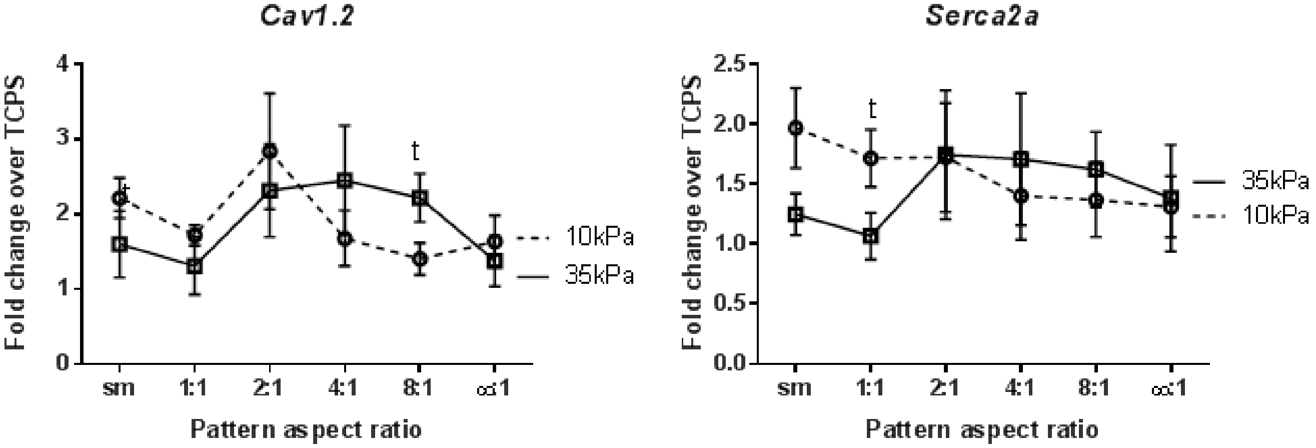
Cav1.2
Fold change over TCPS
4
3
2
1
0
sm
1:1
2:1
4:1
8:1
∞:1
Pattern aspect ratio
t
t
10kPa
35kPa
Serca2a
Fold change over TCPS
2.5
2.0
1.5
1.0
0.5
0.0
sm
1:1
2:1
4:1
8:1
∞:1
Pattern aspect ratio
t
35kPa
10kPa

**Figure S1**

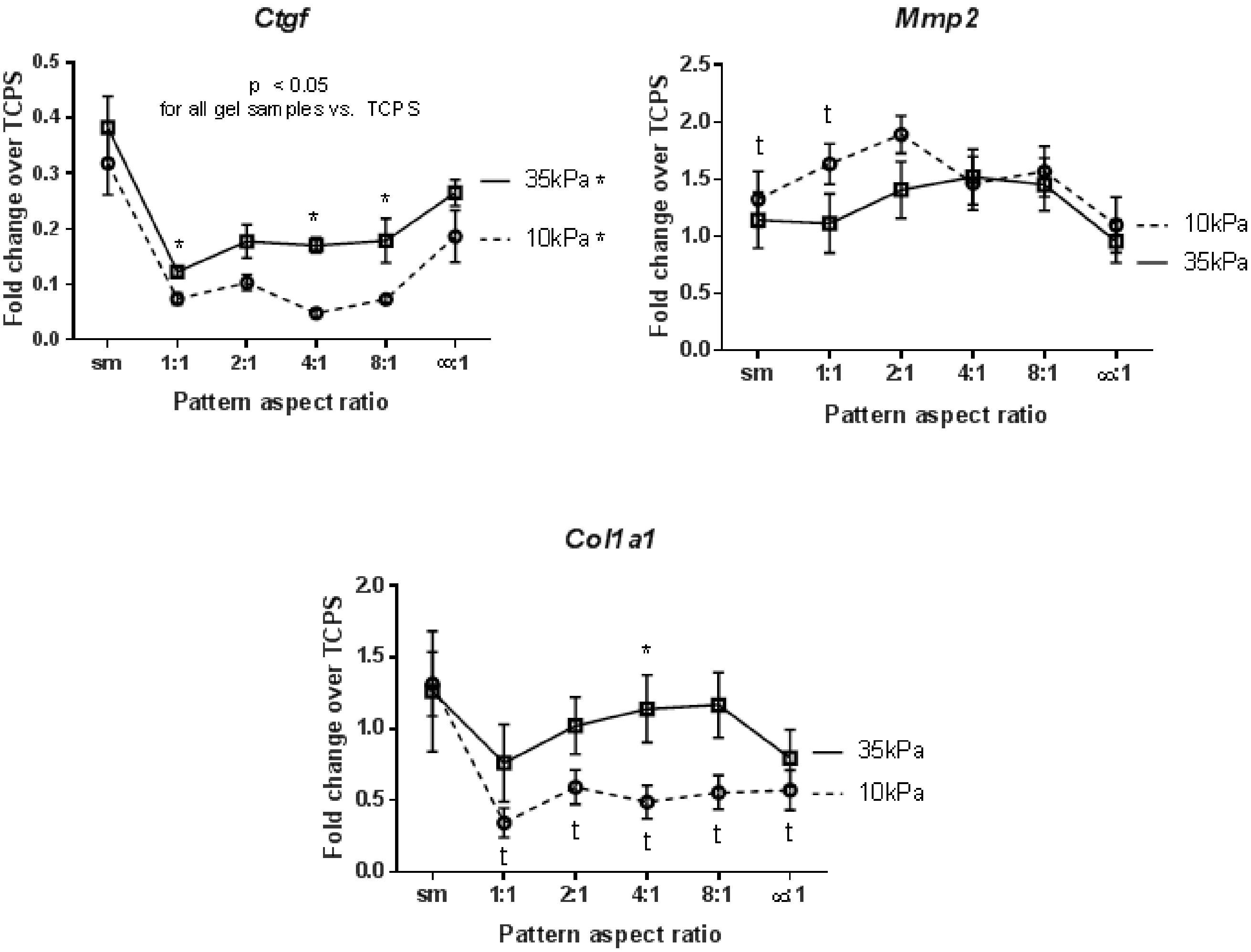
Ctgf
p < 0.05
for all gel samples vs. TCPS
35kPa *
10kPa *
Fold change over TCPS
Pattern aspect ratio
sm
1:1
2:1
4:1
8:1
∞:1
Mmp2
10kPa
35kPa
Fold change over TCPS
Pattern aspect ratio
Col1a1
35kPa
10kPa
Fold change over TCPS
Pattern aspect ratio

Figure S2
